# Inferring protein from mRNA concentrations using convolutional neural networks

**DOI:** 10.1101/2023.11.06.565778

**Authors:** Patrick Maximilian Schwehn, Pascal Falter-Braun

## Abstract

Transcript abundance is a widely used but poor predictor of protein abundance. As proteins are the actual agents executing biological functions, and because signaling outcome depends in a non-linear manner on the concentration of the network components, we aimed to develop a convolutional neural network-(CNN-) based predictor for *Homo sapiens* and the reference plant *Arabidopsis thaliana*. After hyperparameter optimization and initial analysis of the training data, we employed a distinct training module for value and sequence data, respectively, predicting 40% of the variance in protein levels in *Homo sapiens*, respectively 48% in *Arabidopsis thaliana*. Codon counts and peptides had the greatest predictive power. Extracting the learned weight revealed generally similar trends but also some intriguing differences between human and Arabidopsis. Many learned motifs in the 5’ and 3’ UTRs correspond to previously described regulatory features demonstrating that the model can learn ab initio mechanistically relevant features.

## INTRODUCTION

For the predictive analysis of biological systems and the modelling of molecular processes it is essential to know the context-dependent quantitative protein inventory. Essentially all biological processes, including metabolism, signaling, transport, mechanical processes and immune responses are mediated by proteins. Therefore, precise regulation of protein expression is a fundamental process for all organisms to survive in different environmental conditions [1] [2]. However, although gene transcription can be efficiently measured by bulk or single cell sequencing technologies, the resulting protein concentrations are only poorly correlated to transcript levels [3] [4]. At the same time, mere quantitative changes of protein concentrations can have a major impact, e.g., on signal processing and thus cellular behavior. Thus, for accurate analyses and successful modeling of biological systems, it is essential to know the protein concentrations in diverse cell types and disease conditions.

The experimental quantitation of proteomes remains a technically challenging and costly procedure [5]. In contrast, the acquisition of systems-level transcriptomic data is both affordable and common [6]. In the absence of experimental abundance data, analytical and modeling approaches either rely on experimental determination of protein concentrations for the conditions of interest, which is limited by cost and throughput, or use transcript levels to approximate protein concentrations. In silico methods that accurately model protein concentration can be expected to increase the precision of many computational analyses. The complexity of proteostatic regulation prevents mechanism-based modeling approaches [7] [8] [9] [10] [11] [12]. However, the recent immense progress in artificial intelligence and machine learning have enabled the successful development of quantitative predictive models for a variety of challenging biological problems [13] [14] [15] [16] [17] [18] [19].

A prerequisite for machine learning approaches is the availability of suited datasets. Recently, matched transcriptome-proteome datasets became available for representative models of mammals, *H. sapiens* [3] and plants, *A. thaliana* [4]. Here, these datasets are used to develop a machine learning model that more accurately predict protein abundance from transcriptome data than a simple one-to-one transfer as is usually assumed implicitly. Previous efforts to use machine learning to infer protein concentrations used explicitly defined input features of the mRNA, e.g., start- and stop-codon context and codon usage, or of the protein, e.g. linear peptide motifs or hydrophobicity [14] [20]. To make less assumptions that could bias the results and to shorten the trial-and-error process of selecting relevant features here we use convolutional layers that self-learn sequence-based features. We experimentally optimized the CNN architecture and then analyzed the weights of the trained CNN, to understand which sequence features contribute most to the protein-to-RNA ratio (PTR).

## MATERIAL AND METHODS

### Datasets

For *H. sapiens*, expression data was collected from [3] and sequence data was collected from Ensembl [21]. For sequence data, release 83 was used for RNA-seq of the expression dataset as well as training of the CNN. For *A. thaliana*, expression data was collected from [4] and sequence data was collected from Araport11 [22]. For the coding sequence and protein sequence, the original RNA-seq dataset 2016-06 was used, but for the untranslated regions we used the newer and larger release 2022-09-14 (38,624 vs 42,736 entries in the 5’-UTR, 33,026 vs 42,347 entries in the 3’-UTR). All sequence features have been preprocessed by one-hot encoding and zero padding to equalize the input size. For the conversion of FPKM values to TPM values, we used the formula TPM = FPKM / sum (FPKM) * 1E6. Log_2_ transformation for TPM, FPKM and histogram features was done with the formula log_2_(x + 1).

### Machine Learning

All experiments have been done in TensorFlow [23] with the default parameters if not mentioned different. For each experiment we computed 5 independent repeats and a 10-fold cross-fold validation. We used stochastic gradient descendent optimization without momentum. The learning rate for the single input features codon count, nucleotide and peptide count and start/stop codon context was set to 1E-3. For the single input features 5’-UTR, 3’-UTR, CDS and protein sequence as well as all combined feature experiments the learning rate was set to 1E-4. For all single input features experiment we computed 256 epochs and for all combined feature experiments 512 epochs. The batch size was set to 32 for all experiments. As a loss function, we used a custom implemented NaN-MSE since the number of valid datapoints for each gene varies. For the convolutional layers we used 16 filters with a filter size of 8 * 4, 10 * 4 or 12 * 4 for the corresponding 5’-UTR, 3’-UTR and CDS experiments 8nt, 10nt and 12nt. The filter size is determined by the manually selected nucleotide size as well as the one-hot encoding size of 4 for the nucleotide dimensionality. For the protein sequence, the corresponding filter size is 8 * 20, 10 * 20 and 12 * 20 since the one-hot encoding for the amino acids has a dimensionality of 20. For the dense layers we used 32 filters for the first dense layer and 16 filters for the second dense layer in the combined feature experiments and the number of input genes for the TPM experiments.

### Clustering

Clustering the convolutional filters to determine sequence motifs was done in scikit-learn [24]. For each experiment we standardized the convolutional filters independently and applied a cutoff on each channel within the filters where the standard deviation across all nucleotides or amino acids is below or equal to 0.2. Afterwards, we duplicated, padded, and rolled each filter, so that each possible roll combination of each filter was calculated. The result matrix for each feature had the size of (#repeats * #folds * #filters * #filtersize / #onehot-dimension, (2 * #filtersize + 1) * #onehot-dimension) with a one-hot dimension of 4 for nucleotide input features and 20 for amino acid input features. We clustered these matrices using OPTICS [25] with Euclidean metric and an xi value of 1E-2 for nucleotide input features respectively 5E-3 for amino acid input features. After clustering we filtered the clusters from largest to smallest, to ensure that each filter is only considered once, although they have been duplicated and rolled in the beginning. For visualizing, we scaled the clusters to a maximum of 1, centered for the largest peak and cut off nucleotides below or equal to 0.1.

### Gene Ontology

For gene ontology enrichment analysis, we used the webservice Panther [26], the Gene Ontology database from May 2023 [27], the annotation data set ‘GO biological process complete’, Fisher’s Exact test type and False Discovery Rate as correction.

## RESULTS

We used two matched transcriptome-proteome datasets include 29 tissues for *H. sapiens* [3] and 30 tissues for *A. thaliana* [4]. The transcriptome measurements were normalized in fragments per kilobase million (FPKM) for *H. sapiens* and transcript per million (TPM) for *A. thaliana* [28]. Further the transcriptome data was cut off in both datasets for a minimum of 1 FPKM, respectively 1 TPM. To standardize the input data of our machine learning model and avoid inconsistency across samples [29], we transformed the transcriptome measurement of *H. sapiens* into TPM. The proteome measurements were normalized in both datasets in intensity Based Absolute Quantification (iBAQ) [30] with an intensity threshold of 5,000. Both datasets have been collapsed into major isoforms for each splicing variant and log2 transformed.

After transforming, cutting-off and collapsing, we analyzed the distribution of datapoints in both species (**Fig. 1A**). For each organism the genes were grouped by their counts of valid individual or matched datapoints: more or equal than 21 and less than 21 datapoints. To reduce statistical artifacts and increase robustness, we focused our subsequent analyses on those genes with more or equal than 21 matched datapoints, which encompasses 7,606 *H. sapiens* and 11,230 *A. thaliana* genes.

**Figure 1.**
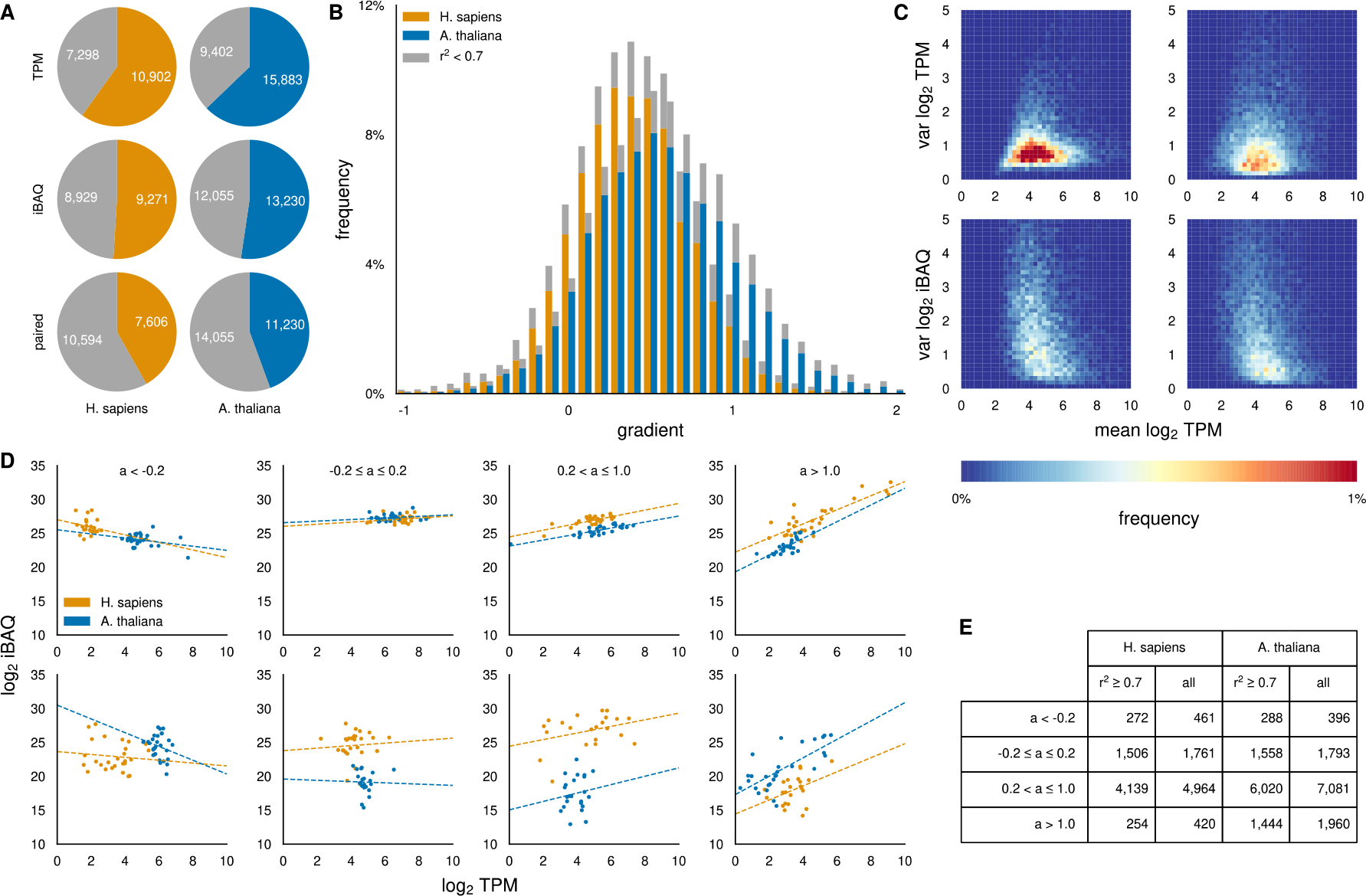
Analysis of both transcriptome-proteome datasets. **A** Gene groups by counts of valid individual or matched datapoints: ≥ 21 (colorized) and < 21 (gray). **B** Distribution of the gradient, poor-quality regressions in gray. **C** Average transcript levels vs. variance of transcript and protein levels (left: *H. sapiens*, right: *A. thaliana*). **D** Representative examples within the indicated gradient bin (top: 100th best, bottom: 100th worst fitting gene). **E** Histogram of the indicated gradient bin.

We first explored which of four commonly used regressors is best suited to describe the relationship between mRNA and protein levels. We applied five independent repeats of a ten-fold cross-validation approach by leaving out tissues and using the unseen data-point pairs to calculate the coefficient of determination (r^2^) (**Ext. Fig. 1A**). Linear regression led to the best fitting predictions and was therefore used as the basis for subsequent analyses. More complex regressors, e.g., squared, performed less robustly, which is often a sign of overfitting due to too few datapoints.

To explore whether linear regression suffers from systematic limitations, we binned genes by the calculated linear gradients and distinguished between good (r^2^ ≥ 0.7) and poor fits (r^2^ < 0.7). The overall distribution of the gradient is gaussian across all genes in both species and without systematic bias in the quality of fits, i.e., good and poor-quality regressions are equally represented across the distribution (**Fig. 1B**). For *H. sapiens*, the distribution of gradients describing the mRNA to protein relationships is narrower, possibly reflecting technical differences in raw data processing, or pointing to differences in the regulation of the translational process in the two species. Surprisingly, in both species we see a proportion of genes that have a negative gradient and a high r^2^. Exploring the dependence of the quality of extrapolation on the number of matched datapoints shows that too few datapoints prevent identification of reasonable relationships, but once at least 21 matched datapoints are available, predictions become reasonably robust (**Ext. Fig. 1B**). Since the cumulative number of matched datapoints sums up from the number of matched datapoints per gene and the frequency of genes within the same bin, leaving out the genes with less matched datapoints does not significantly decrease the available training data (**Ext. Fig. 1C**). In total, we used 213,444 of 256,551 available datapoints for *H. sapiens* and 322,197 of 379,611 available datapoints for *A. thaliana*.

We then explored the effect of average transcript levels on the variance of both transcript and protein levels (**Fig. 1C**). In both modalities and both organisms the greatest variance can be observed at a log_2_ TPM between 3 and 5. While the variance of TPM in H. sapiens is on average lower than the variance of TPM in A. thaliana, the variance of iBAQ in H. sapiens is higher compared to A. thaliana. This indicates that regulation of protein levels in H. sapiens is more likely due to translation regulation, while in A. thaliana transcription regulation seems to have a higher impact on the final protein levels. However, transcripts at lower levels were detectable resulting in a near-gaussian distribution; at lower transcript levels, the corresponding proteins were less readily detected, such that the variance distribution appears truncated.

For illustration, representative examples of genes across the distribution of gradients as well as cumulative numbers are shown in **Figure 1D** and **1E**, respectively. We plotted for both species the 100^th^ best and worst fitting gene within the indicated gradient bin. Clearly, several genes with a negative gradient (a < -0.2) are not statistical artefacts but exhibit a clear trend across many datapoints and with robust coefficients of determination. Nearly a quarter of all analyzed genes have gradients close to zero, indicating that an increase in the mRNA concentration does not substantially impact their protein concentration in the measured steady-state conditions. While ∼85% of these show very robust and constant protein levels, some of the ‘poor fits’ may be non-linear point clouds suggesting complex underlying regulatory mechanisms. Functionally, when focusing on genes with negative gradient (a < - 0.2) and robust good fits (r^2^ ≥ 0,7) we found multiple significant enriched biological processes in both species [26]. In *H. sapiens* the nominal top hits include ‘translation’ (FDR = 2.54E-46, Fisher’s Exact), ‘gene expression’ (FDR = 6.87-31, Fisher’s Exact), ‘metabolic process’ (FDR = 8.55E-24, Fisher’s exact) and ‘biosynthetic process’ (FDR = 1.12E-21, Fisher’s Exact), indicating that these genes are part of core regulatory processes. While in *A. thaliana* the nominal top hits also include ‘protein metabolic process’ (FDR = 4.08E-07, Fisher’s Exact) and ‘macromolecule metabolic process’ (FDR = 9.61E-07, Fisher’s Exact), gene expression related terms are not included but ‘vesicle-mediated transport’ (FDR = 1.09E-06, Fisher’s Exact) and ‘intracellular transport’ (FDR = 6.99E-06, Fisher’s Exact) can be found (**Supplementary Table 1**).To develop a linear regressor for predicting protein-to-RNA ratios, i.e., predicting the gradient (a) and offset (b), we first aimed to identify informative features and thereby investigate the impact of different parts of the mRNA on the protein levels. At the same time, this analysis allowed us to optimize the hyperparameters for each input feature separately. Two different front-end modules were developed to fit different types of input features. For histogram and fixed sequence features, such as codon counts in the actual reading frame and the +1 and +2 frameshift variants, 5’- and 3’-UTR nucleotide counts, amino acid counts, start- and stop-codon context, we used two parallel dense layers combined with a bias operator (**Fig. 2A, top**). The second module was designed for sequence features in sliding windows, which we applied separately to the 5’- and 3’-UTRs, the nucleotide coding sequence (CDS), and the translated peptide sequence. Here, a convolutional layer with 16 filters was combined with a tanh activation and a ReLU activation followed by sum-pooling combined and a log_2_ activation (**Fig. 2A, bottom**). As in the first module we used two parallel dense layers each combined with a bias operator to predict a and b (**Fig. 2A, top**). Finally, we also evaluated all features in a single large network either as such, or after one or two additional dense layers. The predictive power of each sequence feature using this modeling approach as well as the combined sequence feature model after two dense layers are shown in **Figure 2B**.

**Figure 2.**
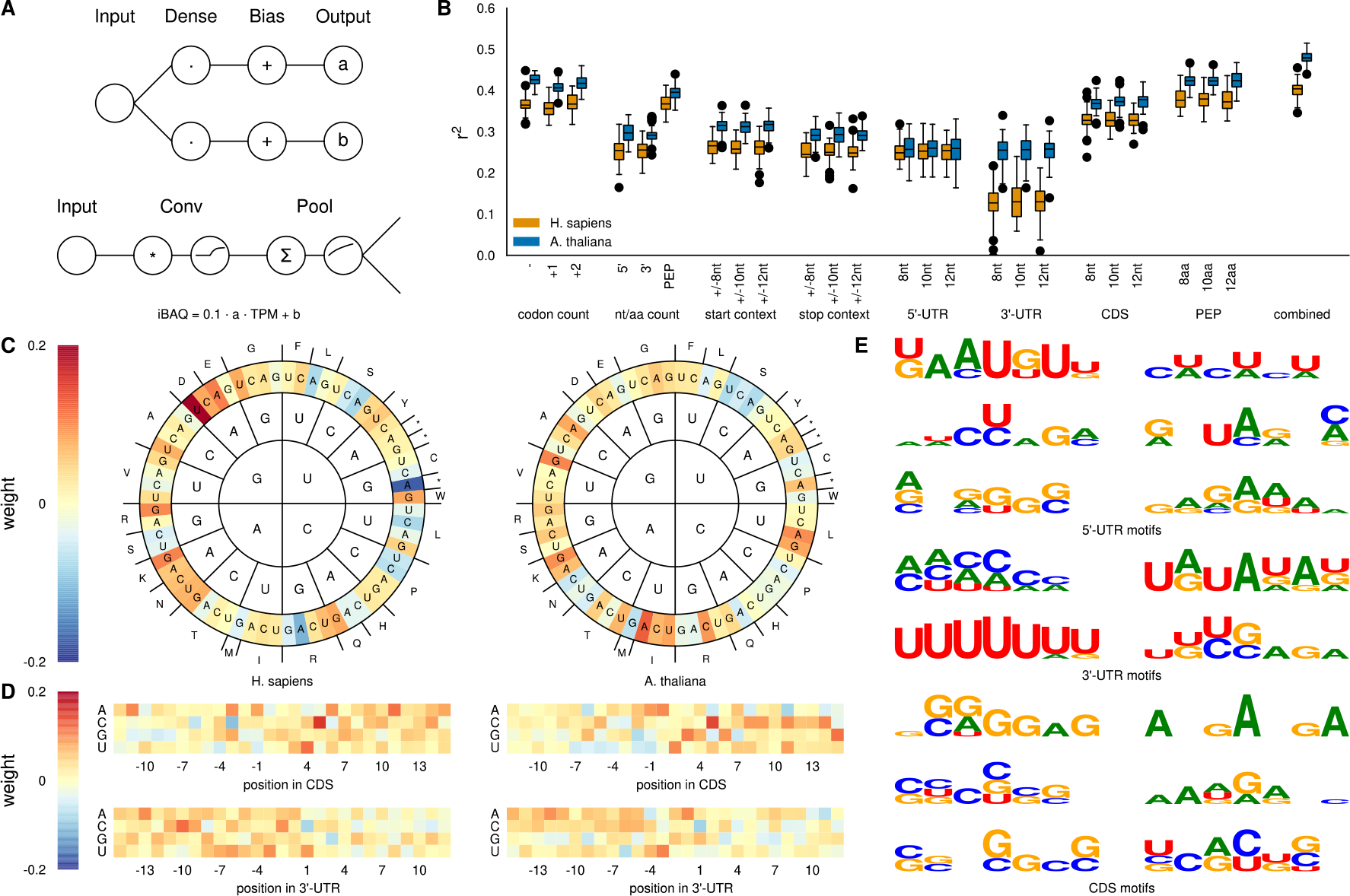
Overview of sequence-based experiments. **A** Circuit diagram of both front-end modules. **B** Predictive power of each input feature across all folds and repeats. **C** Average of learned weights of combined model for codon usage (left: *H. sapiens*, right: *A. thaliana*). **D** Average of learned weights of combined model for start- and stop-codon context (top: start-codon context, bottom: stop-codon context, left: *H. sapiens*, right: *A. thaliana*). **E** Largest clusters of combined model for motifs for each sequence input feature (left: *H. sapiens*, right: *A. thaliana*).

For both species, the coding sequences-derived features (codon counts, amino acid count, CDS and PEP) have the greatest predictive power, with the peptide sequence as the overall best performing single input feature. Among the fixed sequence features, the codon counts are most informative in *A. thaliana*, whereas the corresponding amino acid counts enable slightly better predictions for *H. sapiens*. Somewhat surprising is the observations that out-of-frame codon counts have only slightly lower prediction efficiency than the in-frame counts. It is unlikely that only the nucleotide identity irrespective of the coding potential of each codon impacts translation, considering the good predictive value of the aa counts. More plausible is the hypothesis that even the shifted codon counts reflect underlying trends of the amino acid composition of the full proteins. This can be rationalized by the +2 codon counts, which include the first two bases of the following codon, and, only lacking the often-uninformative wobble position, likely approximate the true amino acid composition.

To explore the details of the codon count, 5’ UTR, and 3’-UTR features we extracted the learned weights of the combined (**Fig. 2C-E**) and the single input models (**Ext. Fig. 2A-E**) without additional dense layer and visualized them as heatmaps by averaging learned weights for the gradient parameter (*a)* across all repeats and folds. Generally, due to the availability of more features to the models, the weights in the combined model are smaller than those in the models trained on each individual feature. Nonetheless, the trends are overall similar but some strong features and motifs that are identified clearly in the individual models, do not emerge or are weaker in the combined model, as some of the respective information is also available from alternative features, e.g., codon count or amino acid count, respectively. Within the single feature experiments, the weights for the overall rare stop codons were significantly higher than the weights of all other codons. We assume that this is without biological context, but a gradient-based learning issue due to overall rare training data for the stop codons. To better display all other codon values, we truncated the color scale at 1. Similarly, we truncated the color scale for the start and stop-codon context to focus on the non-fixed parts of these sequences.

In *H. sapiens*, we observe that a higher count of several codons for the charged amino acids Aspartate (D), Glutamate (E), Lysine (K) and Asparagine (N) have the greatest positive impact on the linear part of the PTR. For Asp (GAC, GAU), Lys (AAA, AAG) and Asn (AAC, AAU) the same effect can be observed when the model is only trained on amino acid composition (**Fig. E2A**). Notably, we observe some similarities but also clear differences in the impact of codons and amino acids on PTR in *A. thaliana*. The codon with the strongest positive impact (AUA) encodes Isoleucine (I), which in plants has a strong positive effect on PTR ratios but not in humans. Other major differences are the strong effect that Asp has in humans but not plants and the differential negative effects of the amino acids Leucine (L) and Serine (S) in both organisms. Generally, we observe that hydrophobic amino acids have a stronger positive weight for PTR prediction in plants than in humans. We speculate that this could be due to the different body temperatures, i.e., 37°C for humans and a highly variable range between 4°C and 34°C for Arabidopsis, which likely impose different constrains on the biophysical demands on proteins, which in turn may have evolved to be reflected in the translation rates.

The weights of the start-codon and stop-codon context for both species reveal that the nucleotides in the CDS have a larger impact on the PTR (**Ext. Fig. 2C**). In the combined model, we additionally see that in both species the strongest impact comes from the second nucleotide (pos 5, both top panels, **Fig. 2D**) after the start codon as well as the first nucleotide after the stop codon (pos 1, both lower panels, **Fig. 2D**). In both cases, the nucleotide C has a strong impact (positive for start-context, negative for stop-context) on the linear part of the PTR.

To analyze the convolutional filters, we clustered the learned weights of each repeat and fold with all possible shift combinations. After clustering each experiment and species separately, we see that in the 5’-UTR of *H. sapiens*, the largest cluster can detect upstream and/or out of frame start codons (AUG) as well as alternative start codons (AUG/CUG) [31] (left, panel 1, **Fig. 2E**). On the contrary, the second largest cluster in the 5’-UTR of *A. thaliana* have been learned to detect stop codons (TAG/TAA) (right, panel 2, **Fig. 2E**). When only using a single input feature, we additionally see in the 5’-UTR of *H. sapiens* the motif UGG (left, panel 3, **Ext. Fig. 2D**), which has been described as tRNA binding site [32] [33]. In the 3’-UTR as well as in the CDS of *H. sapiens*, multiple motifs detect either enriched Uracil content (3’-UTR, left, panel 2, **Fig. 2E**) or enriched Cytosine-Guanin content (CDS, left, panel 2 + 3, **Fig. 2E**). Interestingly, the CDS motifs of *A. thaliana* detect enriched Adenine-Guanin content (right, panel 1 + 2, **Fig. 2E**). Additionally, in the 3’-UTR of *A. thaliana*, the motif UGUA, which can be seen in the combined model (right, panel 1, **Fig. 2E**) and even stronger in the single feature model (right, panel 1, **Ext. Fig. 2D**), has been learned, which is described as binding factor of the cleavage factor I_m_ [34].

Overall, we see a high variance in the predictive power of different features as well as a saturation of the predictive power after creating a combined model. Since the convolutional layers were able to successfully learn motifs, that can be confirmed by literature research, we hypothesized that including the expression score of possible interactors can improve the accuracy further. Therefore, in our next series of experiments we expanded our architecture, to include mRNA expression scores as a proxy of protein expression scores for different manually defined sets of genes (**Fig. 3A**). To manually define the different sets of genes we used two approaches: A quantitative approach by using the linear correlation for each possible gene-gene combination as well as a qualitative approach by selecting genes based on gene ontology terms that are related to either translation or protein decay. Additionally, we used for each set of genes a random selected reference group of the same number of genes.

**Figure 3.**
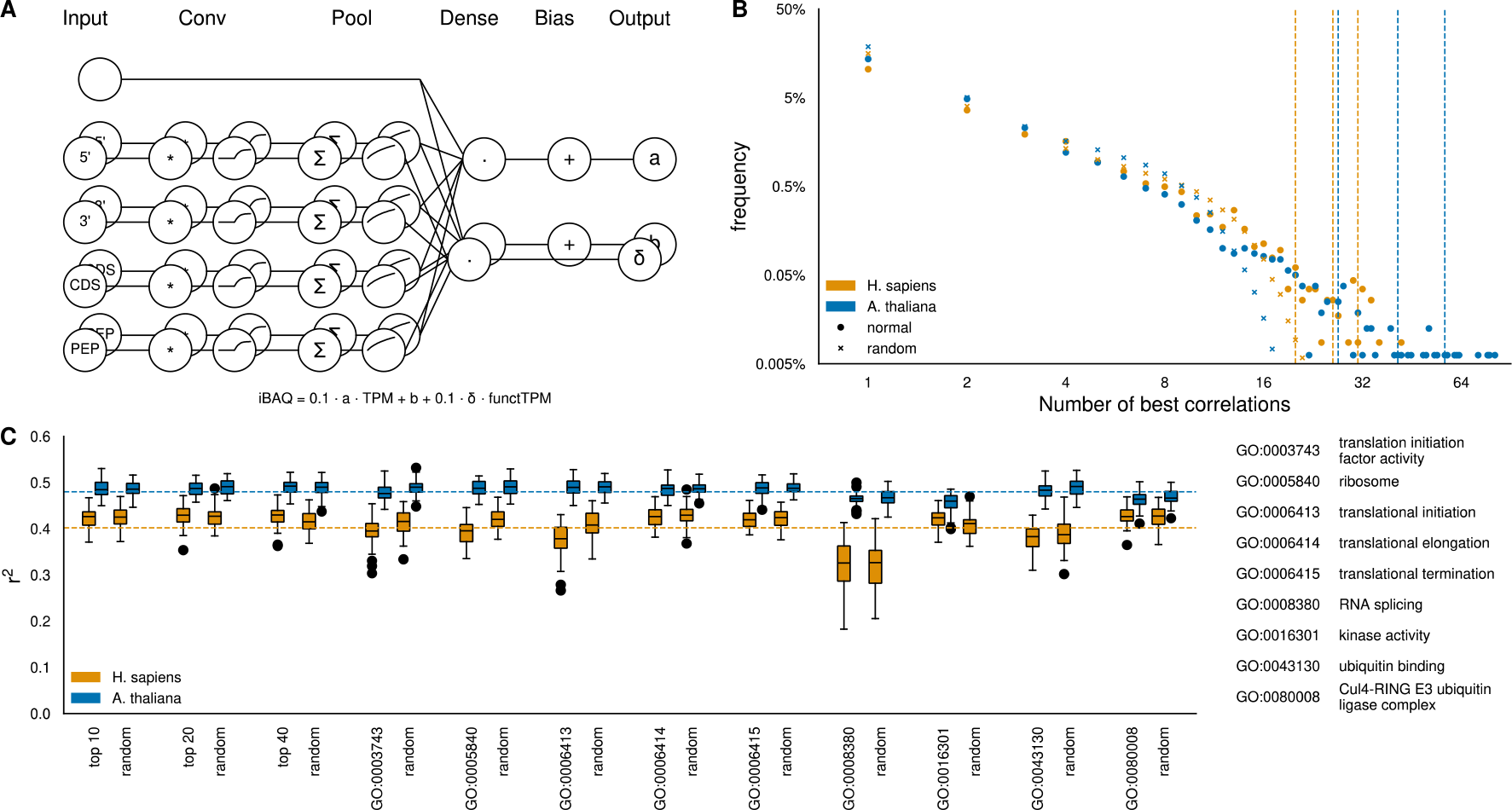
Overview of cell dependent experiments. **A** Expanded circuit diagram. **B** Linear cross-correlation of all gene-gene combinations. **C** Predictive power of each input feature.

For the quantitative approach, we performed a linear regression for each possible gene-gene combination, with cross fold validation and multiple repeats. The result matrix has been collapsed, by selecting for each possible output gene the best fitting linear regression as well as the corresponding input gene. As a result, we see that we have a small number of input genes, which have the highest prediction score of tenth of output genes (**Fig. 3B**). As a result, we see that we have a small number of input genes, which have the highest prediction score of tenth of output genes (**Fig. 3B**). A go enrichment analysis for the top predictors (average r^2^ > 0.8) highlights, that in H. sapiens ‘adaptive immune response’ (FDR = 2.07E-05, Fisher’s Exact), ‘regulation of immune system process’ (FDE = 1.85E-04, Fisher’s Exact) and ‘immune effector process’ (FDR = 2.32E-03, Fisher’s Exact) are the top hits, indicating that immune dependent genes are the key effectors in our dataset. For *A. thaliana*, the top hits include ‘response to stimulus’ (FDR = 7.17E-07, Fisher’s Exact) and especially ‘response to radiation’ (FDR = 1.69E-04, Fisher’s Exact) as well as ‘response to light stimulus’ (FDR = 1.99E-04, Fisher’s Exact), indicating that environmental conditions seem to have the most impact on PTRs of our dataset (**Supplementary Table 2**). We used for each species the top 10, 20 and 40 of the quantitative determined best predictors.

In this series of experiments, we see only a minor impact of the additional input features compared to the best fitting model of previous experiments, often equally with randomly selected genes (**Fig. 3C**). One challenging aspect in learning complex cell-related correlations is that both of our datasets are still limited by their number of tissues (29 for *H. Sapiens*, 30 for *A. thaliana*), while in the first series of experiments, where only sequence dependent features were learned, we can rely on the number of available gene datasets (7,606 for *H. Sapiens*, 11,230 for *A. thaliana*).

## DISCUSSION

In the first series of experiments, we see that the coefficient of determination can be improved stepwise to a r^2^ of 0.40 for *H. sapiens* and 0.48 for *A. thaliana* by adding new features and additional dense layers. By exploring the trained models, we additionally reveal insights which specific input features, e.g., codon, amino acid or short sequence motif, contribute most to the prediction accuracy. In the second set of experiments, analysis of cell-context specific correlations reveals that genes with immune related terms in *H. sapiens* and environmental related terms in *A. thaliana* have the highest correlation to the protein expression levels. Further we see that cell-context dependent approaches are limited by the number of available datapoints for each gene.

Despite the limitation of tissued dependent approaches, our model of tissue independent prediction significantly improves previous applications, by expanding the model to predict the protein-to-RNA ratio for multiple tissues as well as improving the prediction performance. Further improvements of machine learning approaches can be done by including experimental validated protein-protein-interaction data.

**Table 1.**
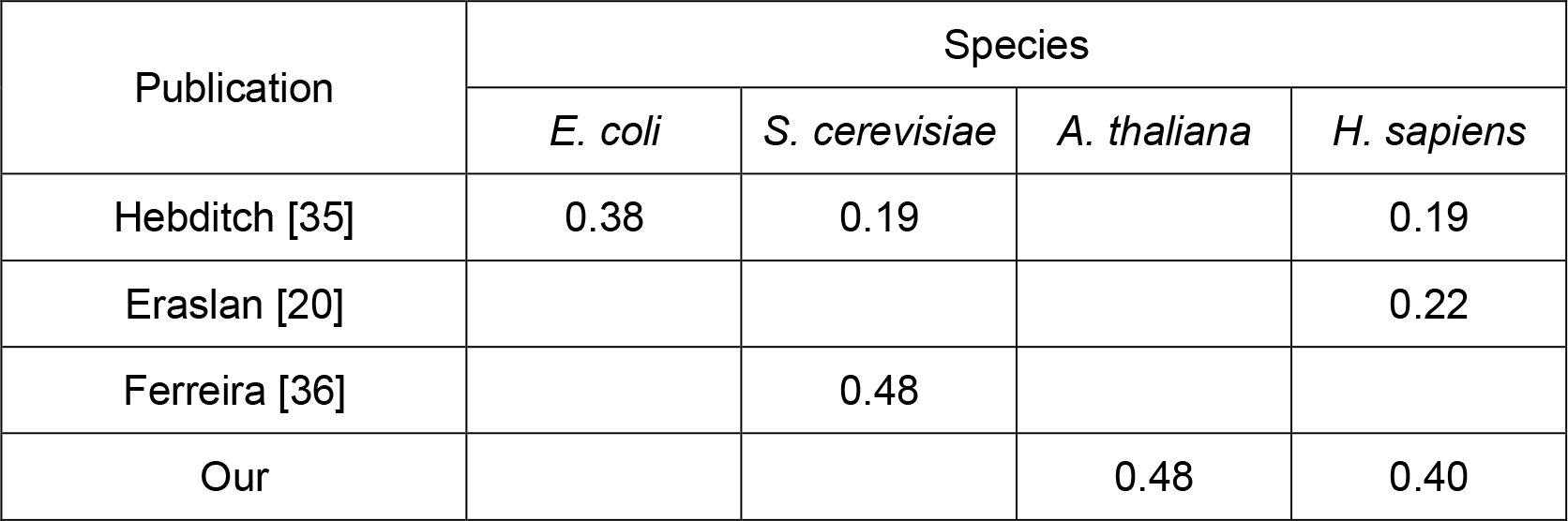
Performance comparison for different publications and species. Performance measured in r^2^.

## FIGURES

**Extended data Figure 1.**
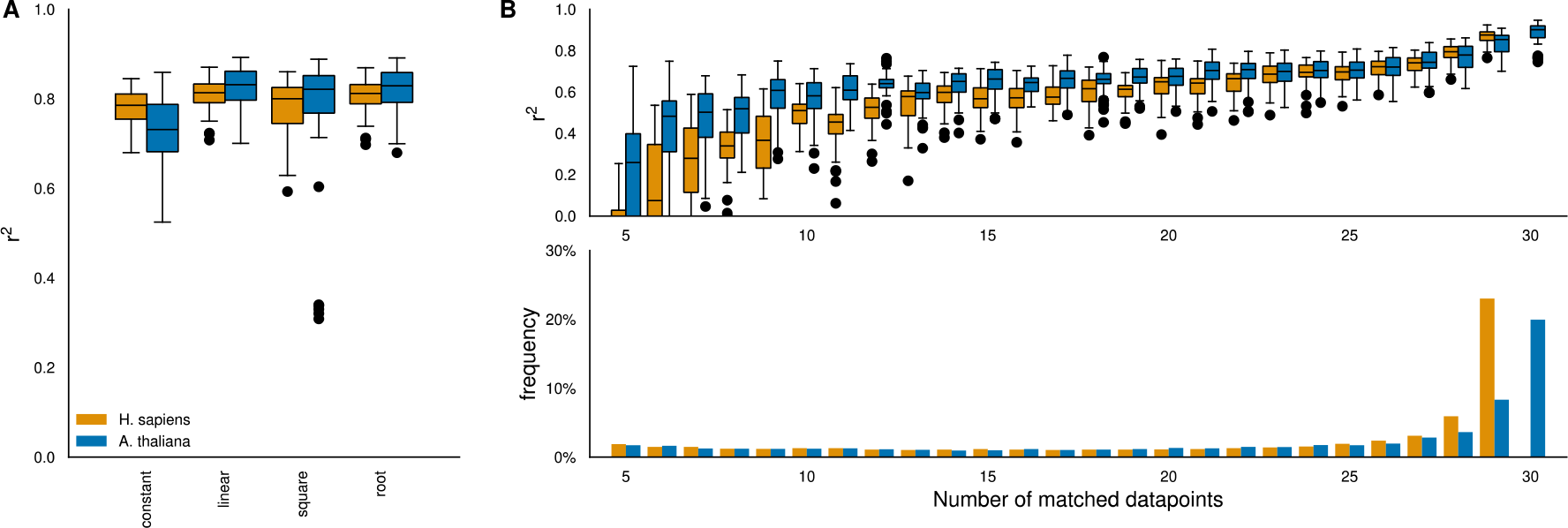
Extended regression analysis. **A** Prediction score of different regressors. **B** Prediction score of linear regressor in correlation with matched datapoints. **C** Frequency distribution of genes and matched datapoints per gene.

**Extended data Figure 2.**
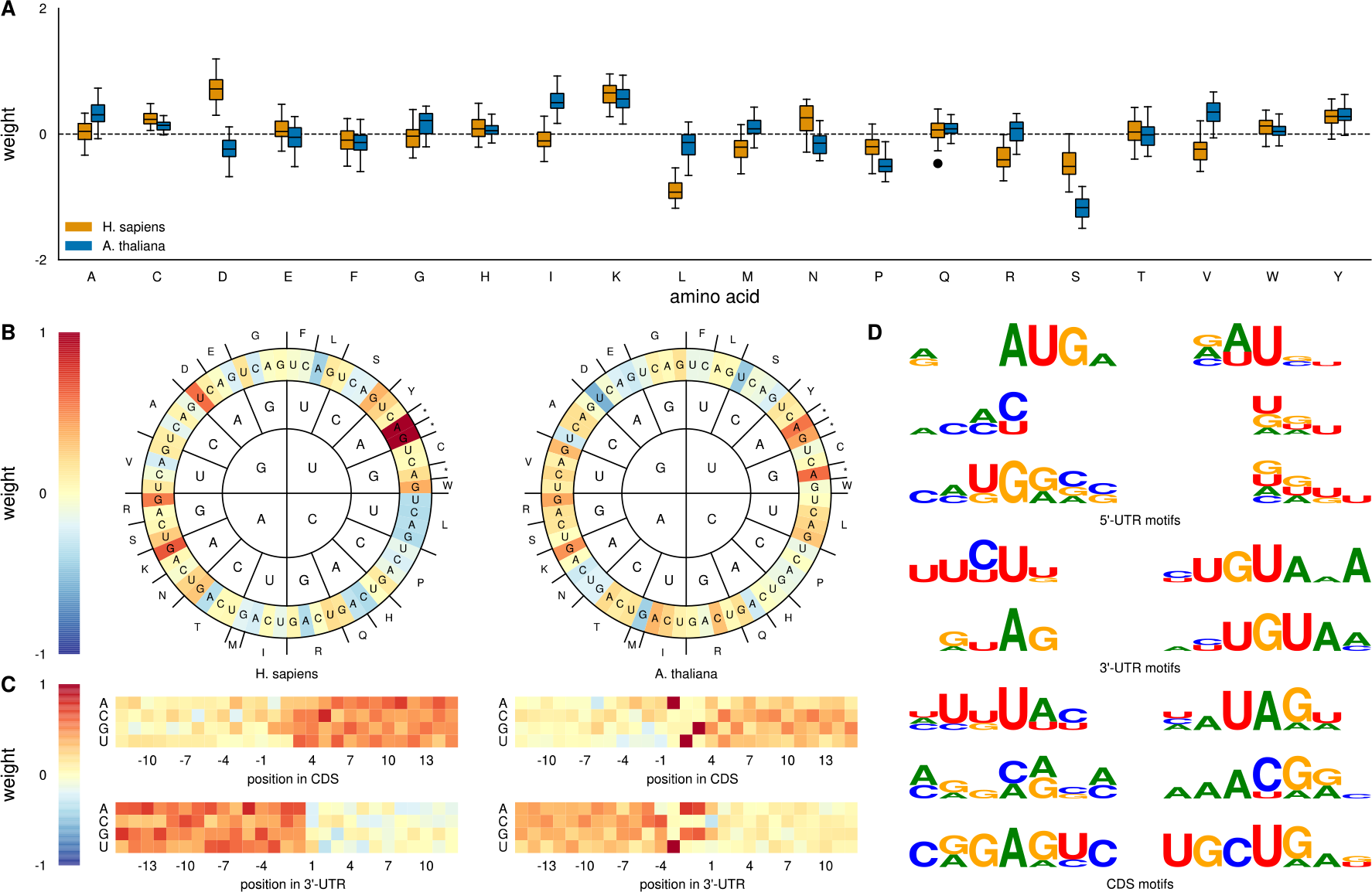
Overview of single input feature experiments. **A** Learned weights of single feature input model for amino acid usage **B** Average of learned weights of single feature input model for codon usage (left: *H. sapiens*, right: *A. thaliana*). **C** Average of learned weights of single feature input model for start- and stop-codon context (top: start-codon context, bottom: stop-codon context, left: *H. sapiens*, right: *A. thaliana*). **D** Largest clusters of single feature input model for motifs for each sequence input feature (left: *H. sapiens*, right: *A. thaliana*).

## TABLES

**Supplementary Table 1** Functional analysis of genes with negative gradient (a < -0.2) and robust good fits (r2 ≥ 0.7). Raw values of linear regressions for **A** *H. sapiens* **B** *A. thaliana*. Gene Ontology enrichment analysis for **C** *H. sapiens* **D** *A. thaliana*.

**Supplementary Table 2** Functional analysis of genes with good average fits (r2 ≥ 0.8). Raw values of linear regressions for **A** *H. sapiens* **B** *A. thaliana*. Gene Ontology enrichment analysis for **C** *H. sapiens* **D** *A. thaliana*.

## ACKNOWLEDGEMENTS

This work was supported by the BMBF-funded de.NBI Cloud within the German Network for Bioinformatics Infrastructure (de.NBI) (031A532B, 031A533A, 031A533B, 031A534A, 031A535A, 031A537A, 031A537B, 031A537C, 031A537D, 031A538A).

## REFERENCES

1. Merchante, C., A.N. Stepanova, and J.M. Alonso, Translation regulation in plants: an interesting past, an exciting present and a promising future. Plant J, 2017. 90(4): p. 628–653.

2. Buccitelli, C. and M. Selbach, mRNAs, proteins and the emerging principles of gene expression control. Nat Rev Genet, 2020. 21(10): p. 630–644.

3. Wang, D., et al., A deep proteome and transcriptome abundance atlas of 29 healthy human tissues. Mol Syst Biol, 2019. 15(2): p. e8503.

4. Mergner, J., et al., Mass-spectrometry-based draft of the Arabidopsis proteome. Nature, 2020. 579(7799): p. 409–414.

5. Aebersold, R. and M. Mann, Mass spectrometry-based proteomics. Nature, 2003. 422(6928): p. 198–207.

6. Wang, Z., M. Gerstein, and M. Snyder, RNA-Seq: a revolutionary tool for transcriptomics. Nat Rev Genet, 2009. 10(1): p. 57–63.

7. Paik, I., S. Yang, and G. Choi, Phytochrome regulates translation of mRNA in the cytosol. Proceedings of the National Academy of Sciences, 2012. 109(4): p. 1335–1340.

8. Shalgi, R., et al., Widespread Regulation of Translation by Elongation Pausing in Heat Shock. Molecular Cell, 2013. 49(3): p. 439–452.

9. Barbosa, C., I. Peixeiro, and L. Romãoa, Gene expression regulation by upstream open reading frames and human disease. PLoS Genet, 2013. 9(8): p. e1003529.

10. Hinnebusch, A.G., I.P. Ivanov, and N. Sonenberg, Translational control by 5’-untranslated regions of eukaryotic mRNAs. Science, 2016. 352(6292): p. 1413–6.

11. Hanson, G. and J. Coller, Codon optimality, bias and usage in translation and mRNA decay. Nat Rev Mol Cell Biol, 2018. 19(1): p. 20–30.

12. Wek, R.C., Role of eIF2alpha Kinases in Translational Control and Adaptation to Cellular Stress. Cold Spring Harb Perspect Biol, 2018. 10(7).

13. Alipanahi, B., et al., Predicting the sequence specificities of DNA- and RNA-binding proteins by deep learning. Nat Biotechnol, 2015. 33(8): p. 831–8.

14. Munusamy, P., et al., De novo computational identification of stress-related sequence motifs and microRNA target sites in untranslated regions of a plant translatome. Sci Rep, 2017. 7(1): p. 43861.

15. Cuperus, J.T., et al., Deep learning of the regulatory grammar of yeast 5’ untranslated regions from 500,000 random sequences. Genome Res, 2017. 27(12): p. 2015–2024.

16. Zhuang, Z., X. Shen, and W. Pan, A simple convolutional neural network for prediction of enhancer-promoter interactions with DNA sequence data. Bioinformatics, 2019. 35(17): p. 2899–2906.

17. Eraslan, G., et al., Deep learning: new computational modelling techniques for genomics. Nat Rev Genet, 2019. 20(7): p. 389–403.

18. Zrimec, J., et al., Deep learning suggests that gene expression is encoded in all parts of a co-evolving interacting gene regulatory structure. Nat Commun, 2020. 11(1): p. 6141.

19. Buric, F., et al., The amino acid sequence determines protein abundance through its conformational stability and reduced synthesis cost. bioRxiv, 2023: p. 2023.10.02.560091.

20. Eraslan, B., et al., Quantification and discovery of sequence determinants of protein-per-mRNA amount in 29 human tissues. Mol Syst Biol, 2019. 15(2): p. e8513.

21. Martin, F.J., et al., Ensembl 2023. Nucleic Acids Res, 2023. 51(D1): p. D933–D941.

22. Cheng, C.Y., et al., Araport11: a complete reannotation of the Arabidopsis thaliana reference genome. Plant J, 2017. 89(4): p. 789–804.

23. TensorFlow-Developers, TensorFlow. Zenodo, 2023.

24. Pedregosa, F., et al., Scikit-learn: Machine Learning in Python. Journal of Machine Learning Research, 2011. 12: p. 2825–2830.

25. Ankerst, M., et al., OPTICS: Ordering points to identify the clustering structure. Sigmod Record, Vol 28, No 2 - June 1999, 1999. 28(2): p. 49–60.

26. Thomas, P.D., et al., PANTHER: Making genome-scale phylogenetics accessible to all. Protein Sci, 2022. 31(1): p. 8–22.

27. Carbon, S. and C. Mungall, Gene Ontology Data Archive. Zenodo, 2023.

28. Conesa, A., et al., A survey of best practices for RNA-seq data analysis. Genome Biol, 2016. 17(1): p. 13.

29. Wagner, G.P., K. Kin, and V.J. Lynch, Measurement of mRNA abundance using RNA-seq data: RPKM measure is inconsistent among samples. Theory Biosci, 2012. 131(4): p. 281–5.

30. Schwanhausser, B., et al., Global quantification of mammalian gene expression control. Nature, 2011. 473(7347): p. 337–42.

31. Kearse, M.G. and J.E. Wilusz, Non-AUG translation: a new start for protein synthesis in eukaryotes. Genes Dev, 2017. 31(17): p. 1717–1731.

32. Citti, C., et al., Spiroplasma citri UGG and UGA tryptophan codons: sequence of the two tryptophanyl-tRNAs and organization of the corresponding genes. J Bacteriol, 1992. 174(20): p. 6471–8.

33. Gamper, H.B., et al., The UGG Isoacceptor of tRNAPro Is Naturally Prone to Frameshifts. Int J Mol Sci, 2015. 16(7): p. 14866–83.

34. Yang, Q., G.M. Gilmartin, and S. Doublie, Structural basis of UGUA recognition by the Nudix protein CFI(m)25 and implications for a regulatory role in mRNA 3’ processing. Proc Natl Acad Sci U S A, 2010. 107(22): p. 10062–7.

35. Hebditch, M., et al., Protein–Sol: a web tool for predicting protein solubility from sequence. Bioinformatics, 2017. 33(19): p. 3098–3100.

36. Ferreira, M., et al., Protein Abundance Prediction Through Machine Learning Methods. Journal of Molecular Biology, 2021. 433(22): p. 167267.

